# Fish harvesting advice under climate change: a risk-equivalent empirical approach

**DOI:** 10.1101/2020.09.09.289207

**Authors:** Daniel E. Duplisea, Marie-Julie Roux, Karen L. Hunter, Jake Rice

## Abstract

The rate of climate change (CC) has accelerated to the point where it now affects the mid-to long-term sustainability of fishing strategies. Therefore, it is important to consider practical and effective ways to incorporate CC into fisheries advice so that the advice can be considered conditioned to CC. We developed a quantitative model to characterise the empirical relationship between a variable affected by climate and fish production. We then used model projections as a foundation for a risk analysis of CC effects on harvesting of Greenland halibut *Reinhardtius hippoglossoides* in the Gulf of St Lawrence, Canada. The risk-based approach quantified a) the relative change in risk of a status quo fishing strategy under various CC scenarios, and b) the change in fishery exploitation rates required to achieve a management objective over a specified time period at a level of risk considered acceptable (risk equivalent fishery exploitation advice). This empirical approach can be used to develop risk-based advice for any other external variable that affects stock production in addition to climate-related variables and it can be applied in most situations where there is an index of stock biomass and fisheries catch. Shifting the focus from process-based understanding of the responses of fish stocks to CC to quantification of how CC-contributed uncertainty can alter the risks associated with different fishing strategies and/or management options, can ensure timely delivery of robust scientific advice for fisheries under non-stationary environmental conditions.

## 2 Introduction

Scientific advice for fisheries management typically comprises three steps: a) evaluating the present state of a fish stock, b) determining the impact of different levels of fishery removals on the stock relative to objectives specified in longer-term policies, and c) providing probabilities of achieving those objectives for each catch level. This approach has many assumptions, one of the most important being that the environment in which the fish stock occurs will be constant, or vary randomly without trend, over the period covered by steps b and c: the environmental conditions that are assumed for the objectives and those encountered during the projections to quantify impacts of removals. Climate change (CC), however, is a directional and non-random process that potentially alters both resource state and productivity and could render unreliable advice for resource management that does not take it into account[1]. CC could affect the likelihood of achieving most objectives (including making some objectives unachievable) through inter-alia changing how a resource like a fish stock responds to pressures like fishing. During the current period of rapid CC [2,3], the consequences of management actions that occur on various temporal scales could be impacted by changes in climate-driven environmental variables that affect fish stock production and the ability to meet stock objectives. In such cases, taking CC impacts into account would change advice and decisions for stock management. It is therefore wise to consider approaches to operationally integrate climate considerations in the formulation of science advice, even in the absence of detailed process-based understanding of climate-fish production dynamics. Stock by stock, science advice that incorporates CC may or may not differ substantially from advice that does not incorporate it. However, as evidence of CC impacts on marine ecosystems continues to accumulate, science advice consistent with the precautionary approach (PA) should consider environmental variation and CC as factors that can change the risks associated with alternative fishing strategies and/or management options.

CC-informed fisheries advice can be provided by adapting mechanistic population models for stock dynamics to include a key climate forcing environmental variable on one or more specific productivity processes in the model. A simulated CC scenario can then be used to drive future stock states with this modelled mechanistic dependence. There are limitations to this approach however, as it is well-established that statistical relationships between individual environmental variables and specific biological processes often break down as new data becomes available[4]. Stock production processes are likely affected by a suite of environmental factors, the effects of which may be non-linear, synergistic or antagonistic, and change quickly with the influence of different environmental factors on the production processes changing as different factors take on values that are typical (modal or median) or atypical (infrequent or extreme)[3].

Interest in these varying but important non-linear and non-stationary relationships is growing, and fostering research in areas like managing for resilience of systems and processes in the face of CC. CC-informed fisheries advice is already being proposed in efforts to improve setting objectives and developing higher level multi-year policy guidance. Nevertheless, however robust the objectives and strategies may be to a non-stationary environment, the expert community must face the task of provision of reliable short-term operational advice relative to those objectives and strategies. If the computations underlying the operational advice assume stationarity of the model parameterisation in relation to stock productivity condition (and they usually do), they are also by default saying that the environment is stable or it is unlikely to impact model parameterisation and associated reference points. Alternatively, they may consider a much larger uncertainty that can encompass non-stationarity within their prediction time frame but this has the impact of making all advice highly uncertain and inevitably leads to foregone yield that may be socio-economically difficult to implement and therefore ignored by decision makers.

By assuming non-stationarity or by casting a very wide uncertainty window to derive science advice are both more vulnerable to error if some of the environmental variation is directional and systematic – as is happening with climate change. If researchers can regularly update their knowledge of these environment-productivity relationships, and the ways that the environment is changing, there may be opportunities to update which environmental factors may be approaching or have passed key thresholds, and are affecting stock productivity. This knowledge can be used to condition risk-based advice based on empirical models parameterized to conditions as local in space and time as information allows. In this sense, we seek an advice conditioning that reflects the overall directional trend in the fish production system forced by plausible CC rather than a strategy that encompasses the full range of possibilities (even implausible ones) or very specific hypotheses about the directional trend. We define risk here as the probability of not achieving a fishery management objective (target) over a specified time period[5], given the level of fishing mortality, uncertainty in the assessment, and we have explicitly highlighted additional uncertainty contributed by CC. Risk equivalency is the process of adjusting the fishing mortality (or any other managed aspects of the fishery) in order to maintain a level of risk considered acceptable to achieve the management objectives considering additional sources of uncertainty. Therefore, risk equivalent advice will ensure comparable probabilities of not achieving a fishery management objective, given additional uncertainty contributed by CC. A full exploration of risk concepts is a field of research in itself and is beyond the scope of this paper. Suffice it to say here, fisheries science generally operates within the scope of the well established PA risk framework within which the main component of risk is the probability of (or not) achieving objectives (such as maintaining stock status above levels below which recruitment may be impaired) and risk equivalency is the expectation that such probability will deliver comparable management outcomes despite external factors acting on the stock.

### 2.1 Fishery modelling approaches that include climate variables

One statistical approach for accounting for CC effects in fisheries assessments is by allowing a particular model parameter to be fitted by a random walk[6] or random effects. A random effect in fisheries models is usually a time-varying autocorrelated process like natural mortality, which does not require a driving mechanistic relationship[7]. However, projecting such models into the future necessarily widens the variance, owing to the lack of an underlying mechanism so the randomness of the parameter “walk” continues multiplicatively into the future. This approach can provide a relatively complete and unbiased accounting of variance because directional processes like CC are subsumed under the aggregation of all possible future environmental combinations underlying the processes in the random walk. However, this can result in the variance window being so large as to make almost all future outcomes plausible. This results in low power to differentiate outcomes of alternative strategies limiting the utility of these models in all but very short term projections necessary for providing tactical advice for fisheries.

An alternative class of models that can be useful is empirical or phenomenological models (EMs). EMs operate at an amalgamated process level, i.e. models which recognise broad amalgamated mechanisms driving a population or biological process, even if the forcing driver may consist of multiple distinct mechanisms with different levels of covariance, non-linearities and threshold effects. EMs recognise past relationships as an indication of future possibilities in an informative but not defining way. Consequently, Ems can provide general rather than the specific predictions of detailed modelling approaches[8]. These modelling approaches should not be viewed in opposition, however, and it is useful to have both. A good example of EMs in CC is that detailed mechanistic global climate models (GCM) driven mostly by physical processes have under-predicted observed sea level rise (SLR)[9–11]. Broader phenomenological models for SLR, called “semi-empirical” models in the CC science community, have made better predictions for SLR than GCM by statistically relating global surface temperature to sea level[12]. These models do not, however, predict SLR at finer regional scales and it is not obvious how to scale them down or if that should even be attempted. EMs are therefore complementary to detailed mechanistic models and use of one approach does not prevent the use of the other. EMs allow rapid development of a broad predictive capacity that may provide the basis for risk mitigation and management such as coastal infrastructure in the case of SLR. Their use could be extended in providing fisheries advice for current needs, providing the same advantages that have been found with semi-empirical models in SLR research.

In fisheries and marine community modelling, EMs have a fairly long history depending on how broadly this class of models is defined. Schaefer’s 1957 biomass surplus production model[13] might be seen as an EM model because it contains only two parameters which subsume many distinct production processes such as individual growth, recruitment and mortality. A variety of size-based empirical models have been used to investigate broad changes in the productivity process of entire communities and size dependence of processes such as predation, growth and mortality[14–19] forming what has become known as size spectrum theory. Size spectrum ideas have been used as macro-ecological methods, to predict community responses to fishing[20], to disentangle species effects while still employing broad parameters like predator-prey size ratios[21,22], and as a basis for exploring balanced harvesting strategies[23]. Although these size based EMs are not usually seen as tactical fisheries models advice for individual species, they provide information as a basis for advice on the impacts of fishing on community structure, more sustainable directions and magnitude for fisheries exploitation, and ecological imbalances that could develop in a marine fish community with continued fishing at different intensities. In the present context of CC effects on fisheries, we propose the EM approach as a central tool in developing timely advice to manage risks associated with management decisions under plausible CC scenarios.

Here we have developed an EM approach as a basis for quantifying the change in risk in fisheries advice and management owing to CC. This advice uses the estimated incremental risk[24] due to climate change to guide directional changes in the amount, location and/or timing of removals, or other aspects of management, which may mitigate that incremental risk, even in the absence of a full understanding of the mechanisms linking the environmental drivers to stock productivity. We detail a case study to illustrate how CC-informed, risk-based advice can be developed in data moderate situations commonly encountered in fisheries. The case involves a cold-water adapted groundfish stock facing increased warm water inflows at the southern limit of its distribution range. We develop and use an EM to provide information on the stock’s likely response to a plausible range of environmental change scenarios, and how these responses would affect the risk of not achieving a fishery management objective and quantify the incremental risk to the stock meeting its conservation objective owing to CC and how risk equivalent advice can be derived accordingly.

## 3 Methods

### 3.1 An EM of fish production dependence on climate

We calculate the surplus production of the stock from the biomass and catch time series[25,26]. The surplus production is a rate of annual net production which is influenced by climate or any other external variable. Surplus production as used here is simply the difference in biomass between two successive years plus the fisheries removals in that year. If the stock grew from one year to the next, the surplus production is positive and if it declined then surplus production is negative. Surplus production can be turned into a specific rate by dividing its value by the biomass (e.g. in the middle of the year). This is equivalent to a population growth rate. We can infer that since these population growth rates remove the impact of fishing (by adding it to biomass difference) these population growth rates reflect changes in stock production owing to natural processes and will reflect changes in stock productivity owing to factors such as climate change and ecosystem changes. We therefore developed an empirical model where the time series of biomass specific surplus production rates are a function of an external variable, we call E:

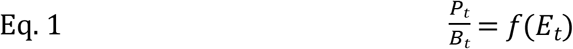

Where *B_t_* is the biomass in year *t* and *P_t_* is the surplus production (*B_t+1_-B_t_*) in year *t* (which is effectively a population growth rate), and *E_t_* is the value of an external (e.g. climate or ecosystem) variable hypothesised to influence surplus production rate. The surplus production rate is like a population growth rate and therefore given an estimate of *B_t_* in a year and a surplus production rate one can determine the production of the stock in that year by multiplying the rate by the biomass and adding it to the previous year’s biomass. This is the simplest kind of surplus production model.

A full derivation of our model is provided in S1 Appendix but the main point is that we derived a simple population growth rate from a time series of survey and catch data and this growth rate is not simply a function of stock characteristics but can be influenced by external variables. For the sake of completeness and testing sensitivity of our analysis we also developed a version with a density dependent term.

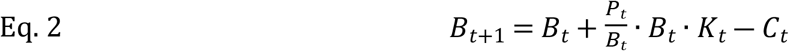

Where *B_t+1_* is the biomass in year *t+1, B_t_* is the biomass in year *t*, *P_t_* is the surplus production in the year *t*, *K_t_* is the density dependent term in year *t* which (*K_t_*≤1) where *K_t_* =1 in the density independent case and *C_t_* is the fishery removals in year *t*. From equation 1 we can see that the surplus production specific rate will be a function of the *E_t_* variable.

### 3.2 Gulf of St Lawrence Greenland halibut (turbot) as a case study

Greenland halibut (*Reinhardtius hippoglossoides*) or turbot is a cold water adapted flatfish with a mean generation time of about seven years. Commercial sized turbot (>40 cm total length) live on the bottom and are concentrated at depths between 100m and 3OOm[27]. The Gulf of St Lawrence (Fig. 1) is located at the southern extent of the species’ distribution range. Recent warming of bottom water in the Gulf[28] is expected to negatively impact turbot production and fishery sustainability. The mechanisms linking warming bottom water temperature and decreased turbot productivity are probably varied and could operate directly through decreases in fecundity, larval and juvenile survival, individual growth and increases in natural mortality of recruited ages, or indirectly through enhanced oxygen depletion in warming bottom waters and associated decreases in individual growth[29].

**Figure 1:**
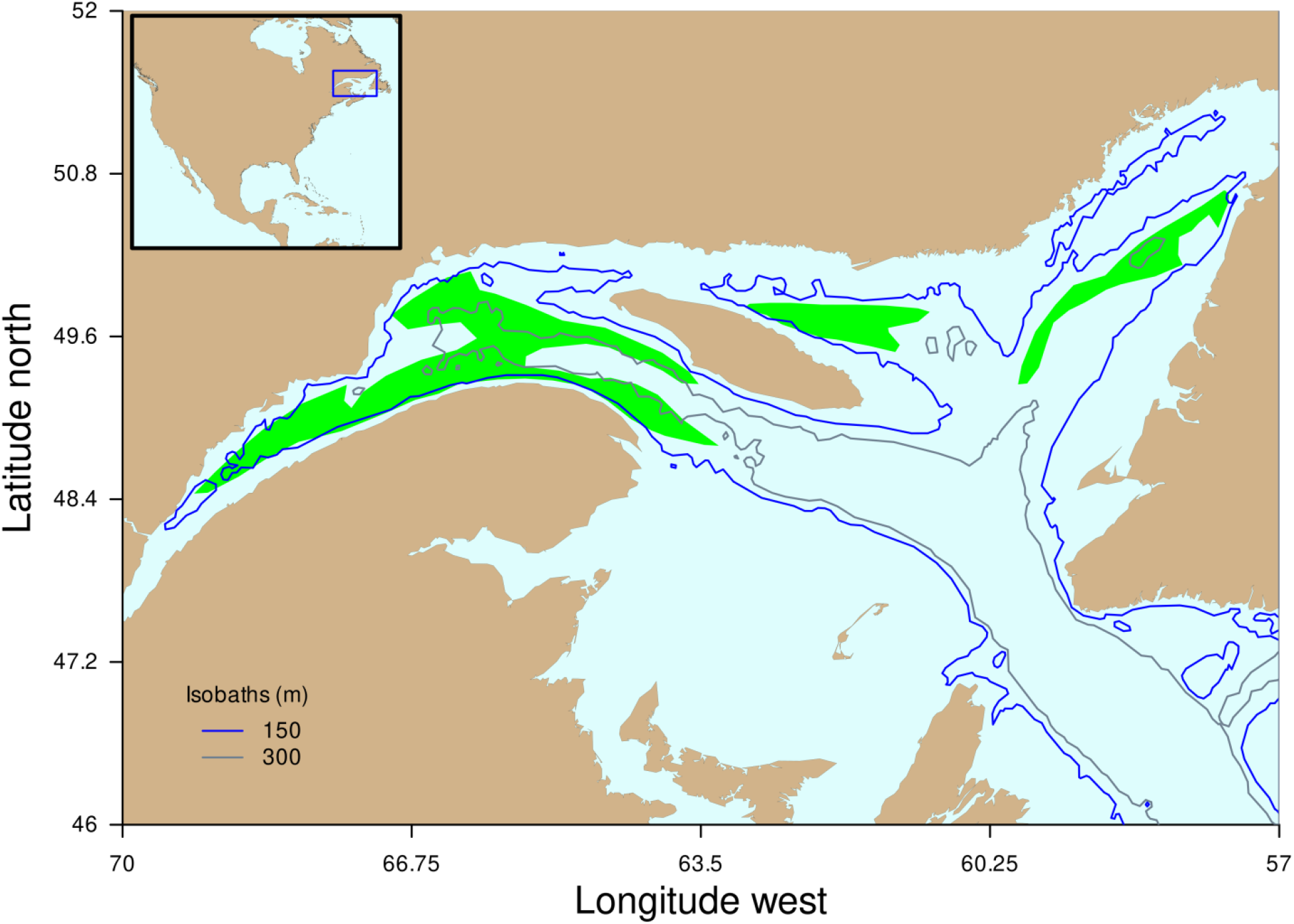
a map of the Gulf of St Lawrence showing the 150m and 300m isobaths and the main turbot fishing areas (in green).

There is a 30-year time series of fishery independent survey catch and environmental data for the turbot stock in the Gulf of St-Lawrence (Fig. 2), but no accepted or benchmarked stock assessment model to assess the impacts of fishing and/or climate on stock dynamics (though one is currently in development). Standard age-based modelling methods have not been used mainly due to age determination challenges. The stock is assessed using biomass indicators primarily derived from fishery independent trawl surveys. Recent science advice for the stock indicated that considering “stock status indicators and ecosystem conditions, a reduction in the exploitation rate may be necessary to promote stock recovery”[30], however no tools are currently in place to quantify how large the reduction should be to have a high likelihood of stock increase.

**Figure 2:**
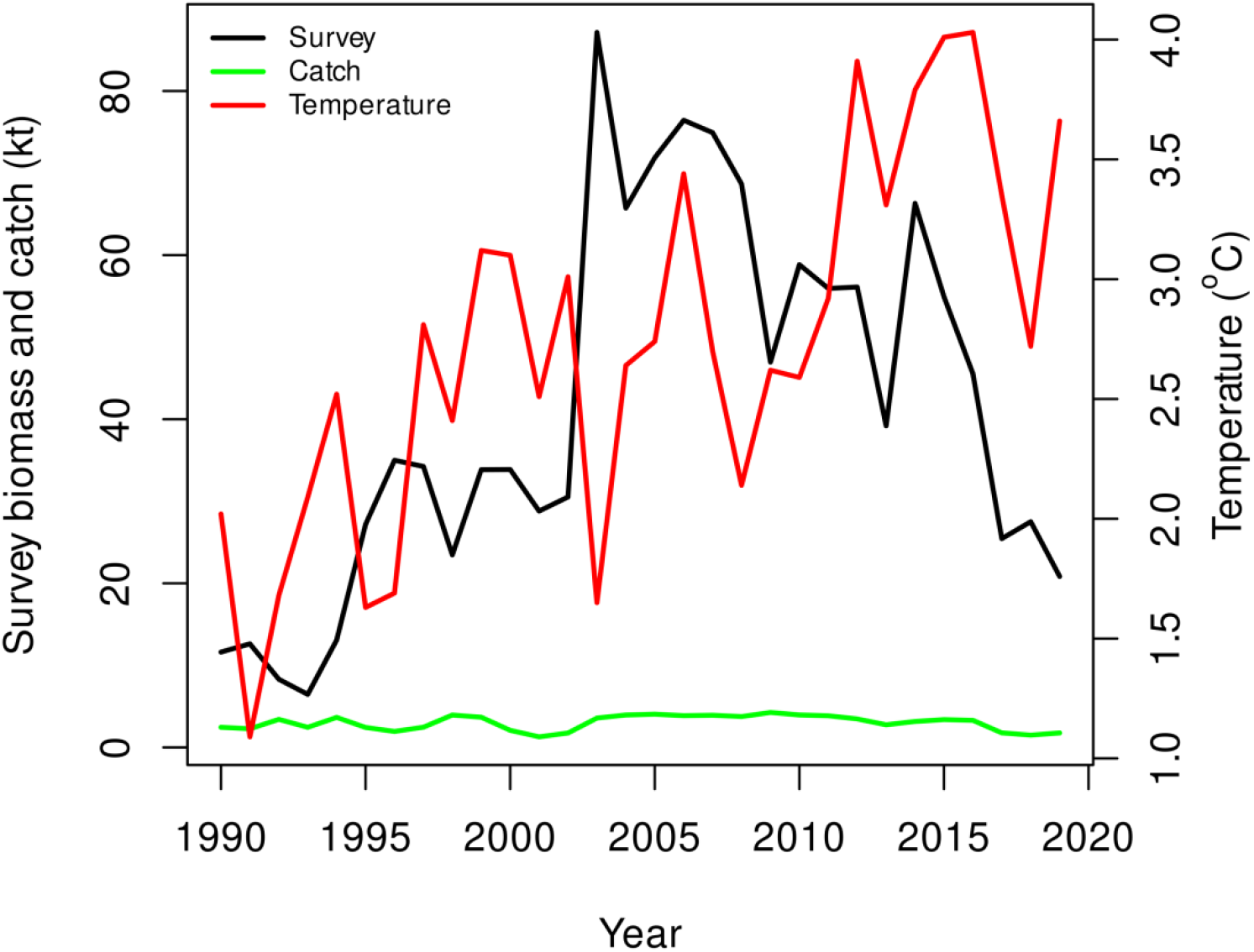
Time series data for Gulf of St Lawrence turbot used to fit the production model. The temperature time series depicted is the annual average temperature of the Central Gulf of St Lawrence water at 150 m.

### 3.3 Model implementation

#### 3.3.1 Production-Environment relationship

*P_t_/B_t_* was calculated from survey biomass and catch time series (fish > 40 cm) over all years which gives n-1 values for biomass surplus production. It is assumed that the catch and biomass index units are the same and of the same magnitude. This assumption would need to be evaluated for any case study in order to put catch and biomass on approximately similar scales but results always remain in relative units regardless. ily derived from fishery independent trawl surveys. Recent science advice for the stock indicated that considering “stock status indicators and ecosystem conditions, a reduction in the exploitAnnual P/B values were statistically related to a time series of annual average water temperature at 150 m in the Central Gulf of St-Lawrence. This temperature index is derived annually based on various observation sources interpolated with bathymetric constraints [28]. Temperature values at depth in the Gulf of St Lawrence do not show large seasonal nor spatial variability, although some mixing still occurs at 150 m in some areas. The temperature at 150 m correlates with a suite of environmental conditions in the Gulf of St Lawrence (e.g. oxygen concentration, temperature at most other, deeper depth strata occupied by Greenland halibut), all of which may be sensitive to CC and impact Greenland halibut production. Thus we refer to temperature at 150 m as the overarching summary variable of CC driven external conditions. A functional relationship between P/B and E was defined using a generalised additive model (GAM). Uncertainty about the relationship was added by resampling the GAM residuals. S2 Appendix provides a more complete explanation of developing the P/B vs E relationship.

#### 3.3.2 Model projections

The P/B vs E relationship formed the basis for projecting the turbot biomass into the future under different fishing climate scenarios. EMs were implemented as per Eq. 2. The density dependent model was implemented under the assumption that the maximum unexploited biomass for the stock was equivalent to 3x the largest index biomass observed. This roughly corresponds to the Bmsy/K value from a Fox surplus production model[31]. Most fish stocks that have been exploited for several generations usually have biomass values at or below Bmsy and therefore we considered the Bmsy/K value of 3 to be realistic and suitable for sensitivity test of the inclusion of a density dependent process. We emphasise that the density dependent version of the model is simply a sensitivity test and the reality is that it makes little difference from the density independent model for a fish stock that has been commercially exploited for many years.

We used the status quo relative exploitation rate in all projections, corresponding to the average exploitation rate observed in the last five data years in our case study (2015-2019). This corresponded to a relative exploitation rate of about 0.05 (i.e. 5% of the population biomass of Greenland halibut >40 cm). This assumption allowed us to investigate how stock production and the probability of achieving a management objective will change under plausible future climate scenarios, if exploitation stays the same.

Climate scenarios were specified by first fitting a normal distribution to E, treating E as a representative of the underlying distribution of observations. Climate conditions were then simulated by shifting the mean of that distribution by ± 0.5 degrees (0.5°C warming and cooling scenarios), and by increasing its standard deviation by a factor of 1.5 (sd x 1.5 scenario) and resampling 20000 times for each year in the future projection. A ‘mean’ climate scenario (corresponding to average observed temperature conditions) was considered, which consisted of re-sampling the distribution fitted to the E series. A ‘null’ model (i.e., model without temperature-dependence on production) was also projected by resampling the observed P/B distribution without regard to the E variable, i.e. the P/B values were simply treated as an empirically derived series and sampled independent of any climate scenario. In all cases but the null model, realisations of future E values were used to determine the population growth rate (P/B) using the GAM model described above and resampled residuals Figure 8: A contour plot showing how exploitation rate of the stock would need to be altered to condition advice for Gulf of St Lawrence turbot to changes in median bottom water temperature at 150 m depth. The climate impact is mediated through the relationship between the temperature and turbot production rate observed in the past. The contours represent the risk expressed as the probability of not achieving the target biomass objective (mean biomass from 1995-2000) after 10 years for different fishing and climate scenarios. For risk levels 50% and lower (shaded green), risk equivalent strategies are possible to compensate for CC by adjusting relative exploitation rate. The actual exploitation rate and temperature scenario is overlaid on the contours.Figure 8: A contour plot showing how exploitation rate of the stock would need to be altered to condition advice for Gulf of St Lawrence turbot to changes in median bottom water temperature at 150 m depth. The climate impact is mediated through the relationship between the temperature and turbot production rate observed in the past. The contours represent the risk expressed as the probability of not achieving the target biomass objective (mean biomass from 1995-2000) after 10 years for different fishing and climate scenarios. For risk levels 50% and lower (shaded green), risk equivalent strategies are possible to compensate for CC by adjusting relative exploitation rate. The actual exploitation rate and temperature scenario is overlaid on the contours.from the GAM model fit. This procedure provided 20000 different projections (Monte Carlo parametric sampling) of the population given the distribution of E (representing plausible future climate scenarios) and status quo F.

#### 3.3.3 Risk analysis and risk-based advice

The risk analysis required specification of three aspects of a healthy stock in a well-managed fishery: (1) a management objective, such as a biomass target from the observed range of biomass values (e.g., B during a given time period); (2) the time by which the management objective must be achieved; and (3) the risk tolerance which is the acceptable probability level for not achieving the objective. The management objective for turbot was established based on a period of relative stability in the survey biomass index and catches from 1995-2000 (Fig. 2). The average biomass (B) (average of annual survey biomass values) over this “baseline” period was used as the biomass target or management objective for the stock. The relatively stable catches during this period (Fig. 2) suggests that the selected biomass objective may have characteristics of a biomass at maximum sustainable yield which is a common management target (DFO 2006). The time period required to achieve the objective was set at 10 years and the maximum acceptable risk of failure to achieve the target within that time was set at 50% (which is the *de facto* risk tolerance for a target).

The annual probability of achieving the objective over the 10-year period given status quo fishing and plausible climate scenarios was determined from the outputs of EM projections, as the proportion of the Monte Carlo projections where the objective was achieved annually or at the end of the 10-year projection. For a given climate scenario, the ratio of the calculated risk under CC forcing relative to the same risk in the null model assuming no productivity dependence on the environmental variable is the incremental risk resulting from consideration of CC effects on stock dynamics:

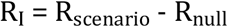

Where incremental risk (R_I_) is the risk under the climate change scenario (R_scenario_) minus the risk under a null model where climate was not considered (R_null_).

For each projection scenario, we can also determine the fisheries exploitation rate that will allow the objective to be achieved at the end of 10 years at the acceptable risk level (50% in the case with a target). The ratio of this exploitation rate in the null model to that in any of the CC scenario is the risk equivalent CC adjustment factor (multiplicative) that would need to be applied to exploitation rate or catch to compensate for CC, we term this the climate change conditioning factor (CCCF):

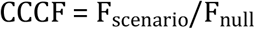

### 3.4 An R package to develop risk equivalent advice under CC

We developed an R package Climate Change Conditioned Advice (ccca) to provide easy access to this method and for transparency (www.github.com/duplisea/ccca). Full instructions for loading the package and running examples with datasets included in the package are provided. This package is easily extendable to any environmental or ecosystem variable (E) for which a time series is available and hypothesised to affect fish production. Different vignettes are also available.

## 4 Results

### 4.1 Stock production-E variable relationship

The relationship between turbot surplus production (production to biomass ratio: P/B) and the E variable was variable, but showed a dome-shaped relationship where P/B was more negative at both extremes of the temperature range and peaked in the middle (Fig. 3). P/B values themselves ranged from about −0.5 to 2 and there was important inter-annual variation in stock production. The P/B vs temperature relationship formed the basis for projecting the turbot biomass into the future under various climate scenarios under the assumption of status quo fishing. Alternative P/B vs E models can be fitted that may be more appropriate representations of hypotheses on external variable influences on production for different stocks (See additional material in https://github.com/duplisea/ccca).

**Figure 3:**
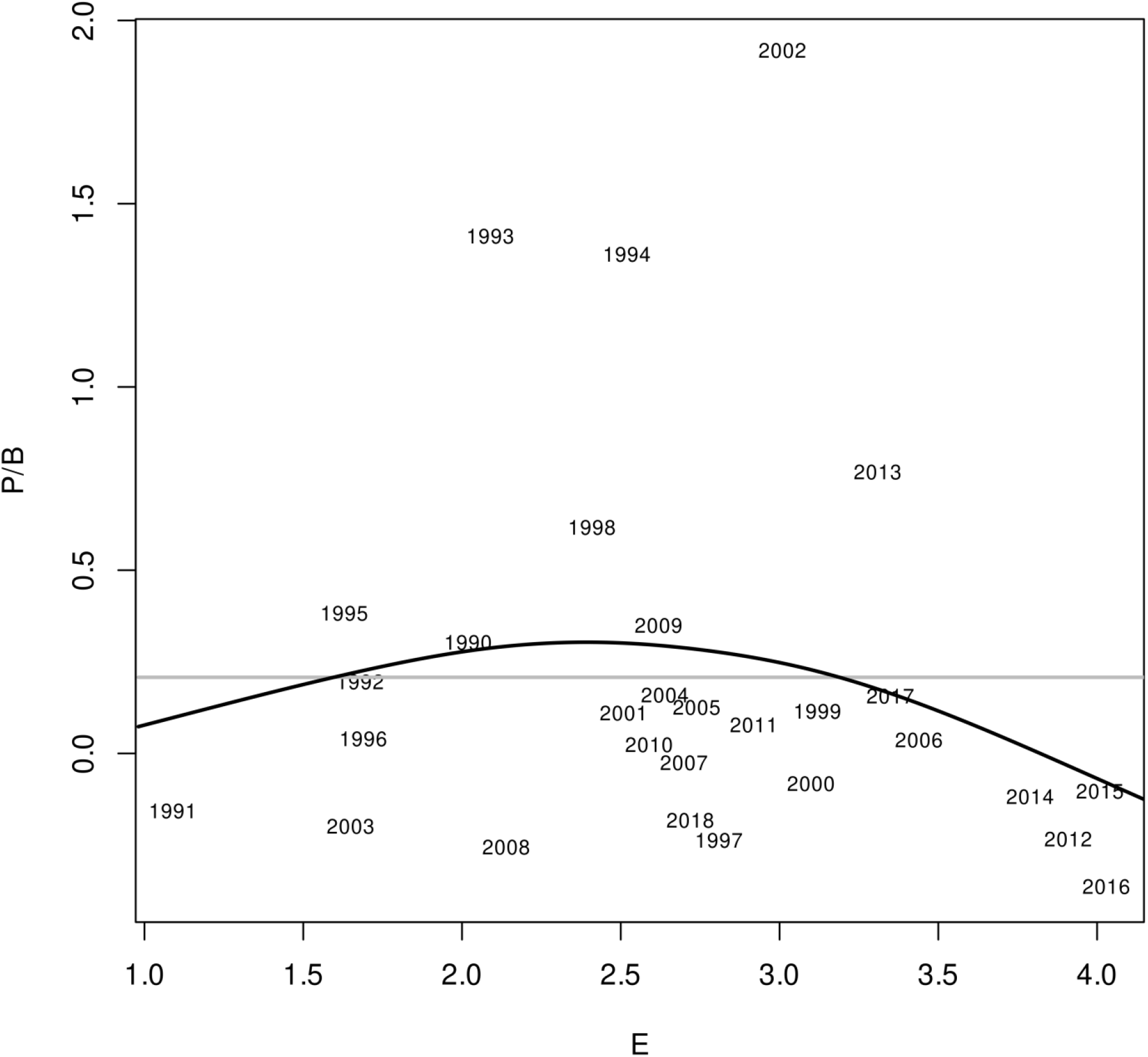
The relationship between the production to biomass ratio (P/B) of Gulf of St Lawrence turbot and the environmental index (annual average temperature in the Gulf ablack) and the null model (grey) is where the E variable has no impact on the P/B aFigure 8: A contour plot showing how exploitation rate of the stock would need to be altered to condition advice for Gulf of St Lawrence turbot to changes in median bottom water temperature at 150 m depth. The climate impact is mediated through the relationship between the temperature and turbot production rate observed in the past. The contours represent the risk expressed as the probability of not achieving the target biomass objective (mean biomass from 1995-2000) after 10 years for different fishing and climate scenarios. For risk levels 50% and lower (shaded green), risk equivalent strategies are possible to compensate for CC by adjusting relative exploitation rate. The actual exploitation rate and temperature scenario is overlaid on the contours.t 150 m depth). A generalised additive model smooth is shown fitted to the data series.

### 4.2 EM projections

The distribution of the four temperature (climate) scenarios used in model projections are shown in Fig. 4. These were used to project future P/B distributions under each temperature scenario (Fig. 5). The climate scenarios represent plausible changes in temperature at 150 m in the central Gulf (0.5 °C warming or cooling) which can be important changes for Greenland halibut in the Gulf of St Lawrence. We also remind the reader that the hypothesis is not that temperature in the central Gulf at 150 m is what the bulk of the fish are experiencing directly but that this this temperature measure has a larger systemic link to Greenland halibut productivity at several depths and locations in the Gulf, i.e. it is something of a metavariable for climate change driven productivity changes pertinent to this stock. The resulting P/B distributions for the stock under each future climate scenario exhibit small but important differences (Fig. 5). The probability density function for P/B is more peaked at values of −0.5 for both the warming scenario and increased variance scenario than for the status quo temperature or cooling scenario. This greater density in the lower part of the P/B range will render projections based on sampling the warmer or more variable pdfs much more pessimistic.

**Figure 4:**
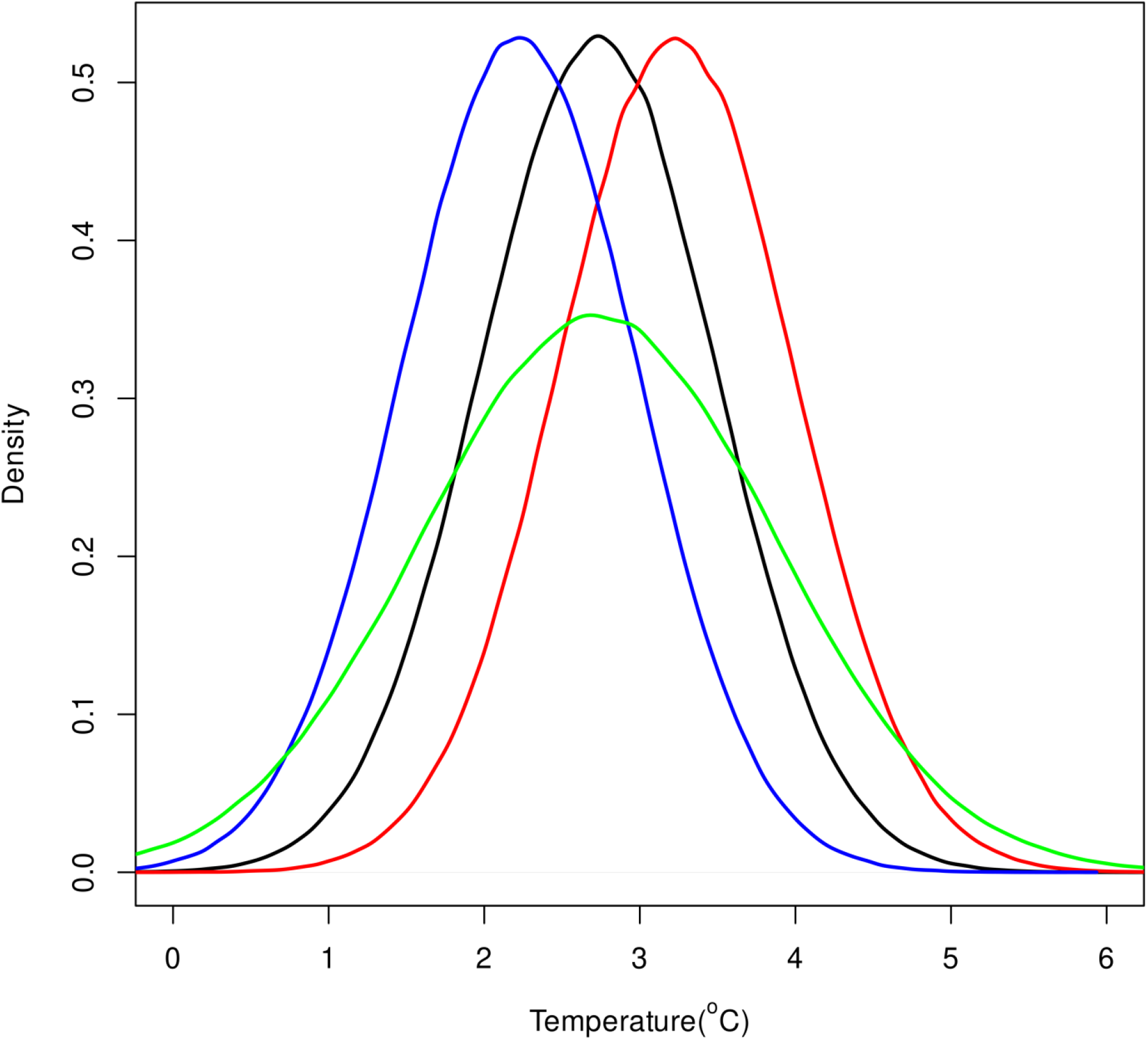
Four temperature scenarios for the environmental variable in the Gulf of St Lawrence over the next 10 years. The black line is status quo, i.e. the future is simply sampled from the past temperature distribution. The red line is the status quo temperature distribution with mean shift 0.5 °C warmer, the blue line the status quo temperature distribution shifted 0.5 °C colder and the green line is the status quo mean but with the standard deviation of the distribution increased by a factor of 1.5.

**Figure 5:**
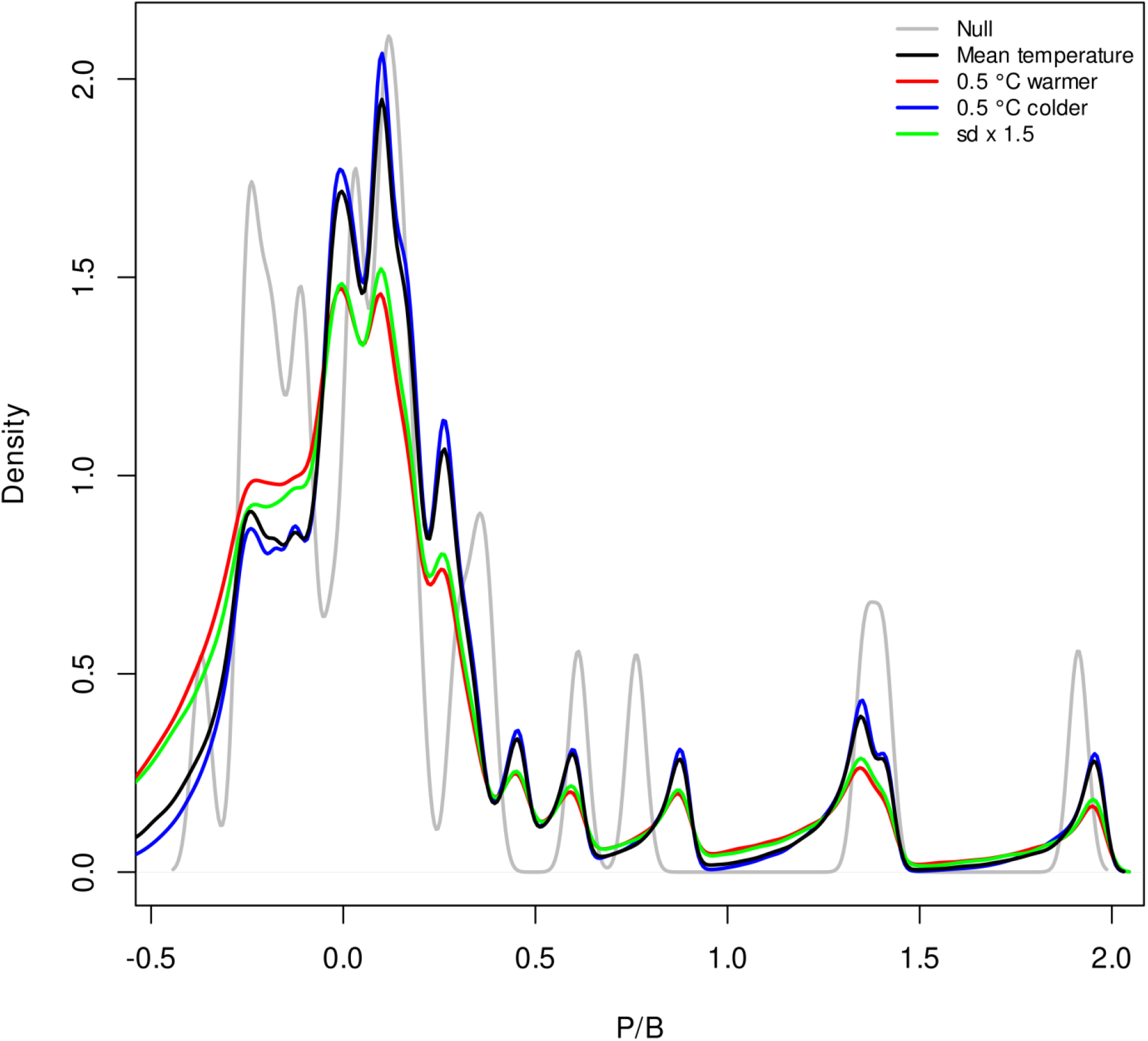
The projected production to biomass ratio (P/B) distribution of Gulf of St Lawrence given the four temperature scenarios (E) and generalised additive model model fit of P/B vs E and also by adding a random residual of that P/B vs E relationship to each future draw. The black line is status quo, i.e. the future is simply sampled from the past temperature distribution. The red line is the status quo temperature distribution with mean shift 0.5 °C warmer, the blue line the status quo temperature distribution shifted 0.5 °C colder and the green line is the status quo mean but with the standard deviation of the distribution increased by a factor of 1.5. The null model P/B (grey) is just the P/B distribution sampled without regard to the E variable.

### 4.3 Risk analysis

The recent mean temperature and 0.5 °C colder scenarios achieved the biomass objective with 50% probability under status quo fishing pressure (Fig. 6). Neither the 0.5 °C warming scenario nor the increased temperature variance scenario allowed the biomass objective to be achieved with 50% probability under status quo levels of fishing. In all scenarios, there was little difference in the ability to achieve the biomass objective (annual probabilities of being at or above the objective) between projections assuming no density dependence vs projections assuming density-dependent stock dynamics. The incremental risk resulting from plausible CC scenarios and risk-equivalent exploitation rates are summarised in Table 1. There was a minimal decrease in risk (1%) when production-dependence on temperature was considered (mean temperature scenario) relative to the null model without temperature-dependence.

**Figure 6:**
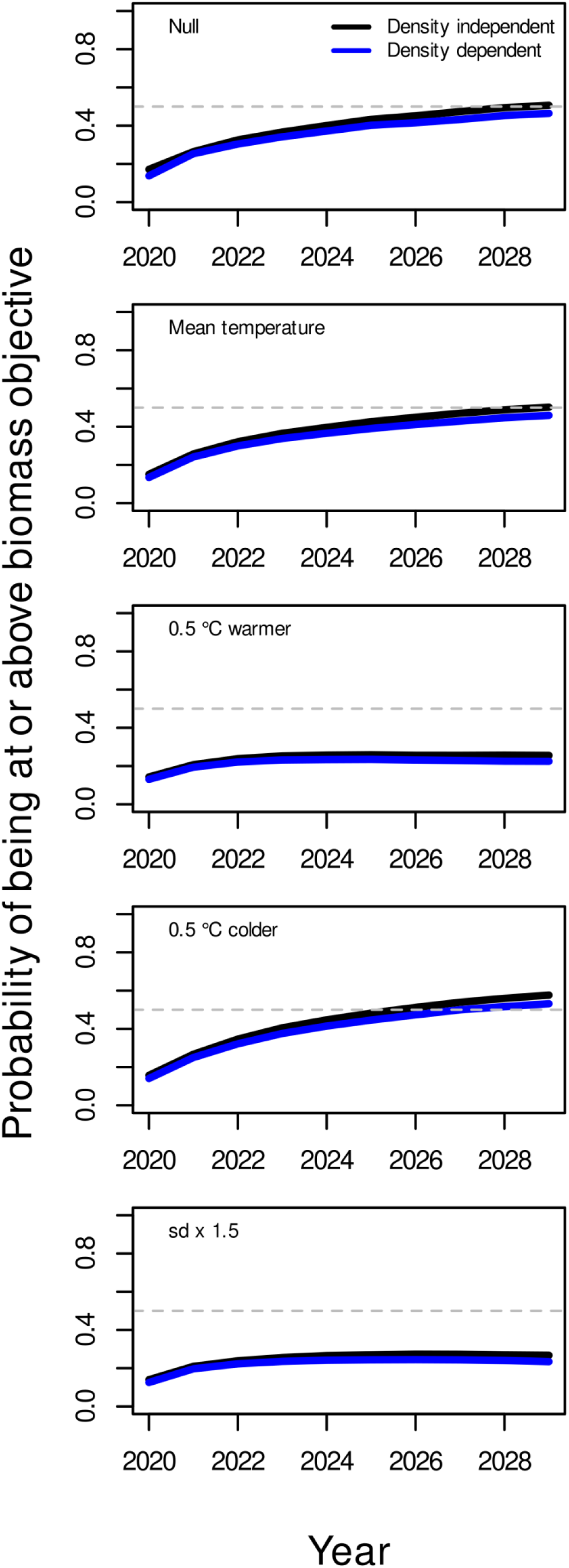
The probability of biomass being at or above its objective (mean biomass from 1995-2000) under each year of a projected climate scenario (2019-2028). The 50% risk or probability is shown as the horizontal dashed line which represents the de facto risk level for a target objective. The probabilities were calculated for both a density independent and density dependent models to determine the impact of the density dependent assumption on risk. A status quo fishing strategy (mean exploitation rate from 2014-2018) was applied for all scenarios.

**Table 1.** Calculated relative risk (1-the probability of achieving the biomass objective at the end of the 10-year projection period (2027)) across scenarios, and incremental risk corresponding to the risk ratio in mean temperature and CC scenarios (numerator) relative to the null model (no dependence on temperature in projections). Risk and incremental risk are calculated under a status quo exploitation rate (5%). NA=Not applicable, NP=Not possible/unachievable.

| CC scenario | Risk (2027) |  | Incremental Risk*<br>(Risk<sub>scenario</sub>-<br>Risk<sub>null</sub>) |  | Risk equivalent exploitation rate<br>(@ 0.50 risk threshold) |  | Climate change conditioning factor<br>(CCCF) |  |
| --- | --- | --- | --- | --- | --- | --- | --- | --- |
|  | Density-independence | Density-dependence | Density-independence | Density-dependence | Density-independence | Density-dependence | Density-independence | Density-dependence |
| Null | 0.494 | 0.536 | NA | NA | 0.071 | 0.058 | 1 | 1 |
| Mean temperature | 0.498 | 0.541 | 0.004 | 0.005 | 0.069 | 0.057 | 0.982 | 0.971 |
| 0.5oC warmer | 0.744 | 0.775 | 0.251 | 0.240 | NP | NP | 0 | 0 |
| 0.5oC colder | 0.423 | 0.468 | -0.070 | -0.067 | 0.089 | 0.077 | 1.269 | 1.324 |
| High variance | 0.733 | 0.766 | 0.239 | 0.230 | NP | NP | 0 | 0 |

The risk increased by 50% (from about 0.5 to 0.75) in the warming scenario compared to the null model and was similar with or without density-dependence. In cooling scenarios, the risk decreased by about 14% relative to the null model. The scenario where temperature variance was increased while the mean temperature remained the same was almost identical to the warming scenario, i.e. the incremental risk to not achieving the stock objective while fishing at status quo increased by 50% (Table 1). For both the warming and increased variance scenarios, the target was unattainable with any level of fishing given the 50% risk standard associated with a target objective (Fig. 7)

**Figure 7:**
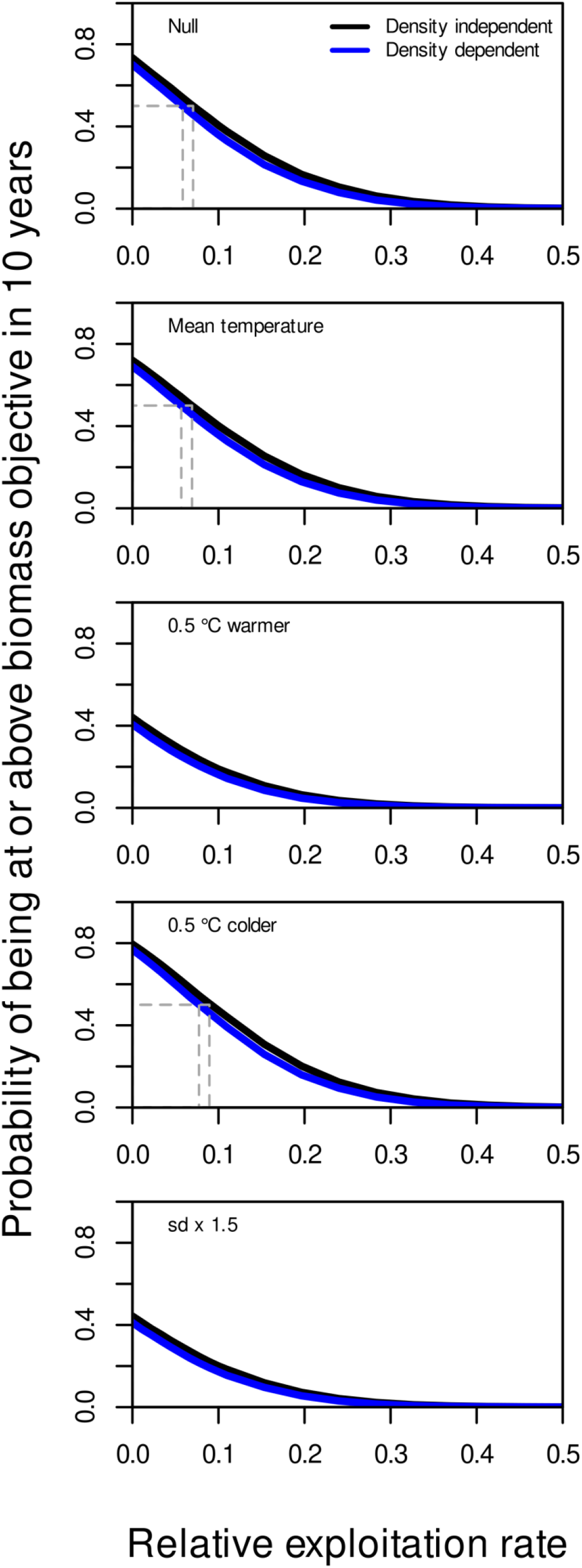
The probability of achieving or exceeding the biomass objective (mean biomass from 1995-2000) after 10 years of a specific climate scenario. The exploitation rate at the 50% risk level (a target) is shown for both density independent and density dependent models.

Adopting a risk tolerance of 50% indicates that the maximum exploitation rate that would allow the biomass objective to be achieved under average temperature conditions (mean temperature scenario 1990-2019) or where productivity had not dependence on temperature (Null) was about 7% and 6% under a density independent and density-dependent assumptions, respectively (Fig. 7, Table 1). For the 0.5 °C colder scenario, these values were about 9% and 8%, respectively and this translates to a climate conditioning factor greater than 1 (the factor to multiply the exploitation rate in order to achieve risk equivalency i.e., maintain the same level of risk associated with the advice) and equivalent to about 1.3 for both density dependent and independent models. Under the 0.5 °C warmer and increased temperature variance scenarios, the biomass objective was not possible to achieve (Table 1 - “NP”) thereby corresponding to a CCCF equal to 0.0 (no exploitation). This does not mean that the objective would be achievable in the absence of exploitation. Rather, it means that there are no other options for management actions in terms of fishery removals that would allow the stock to achieve the objective.

The scope for changing the exploitation rate in relation to CC effects on stock production can be captured by risk equivalent contours for alternative fishing strategies (Fig. 8). The case study here uses a maximum risk tolerance of 50% for the biomass objective. Therefore any risk contour smaller than 0.5 falls within the acceptable risk space associated with the objective (highlighted as the green area in Fig 8). The corresponding median temperature range over this acceptable risk space was about 1.5°C-3.1°C. At about 2.5 °C, a maximal exploitation rate of about 0.09 (or 9% of the population biomass of fish >40 cm), would achieve the biomass objective with a 50% probability over ten years. If median temperature increased to about 3 °C (i.e. a 0.5 °C increase from optimal), an exploitation rate of only about 3% would allow the objective to be achieved in 10 years at a 50% risk level. At median temperatures <1.5 °C and just above 3.1 °C it becomes impossible to achieve the biomass objective in 10 years with a 50% risk tolerance simply by adjusting the fishery exploitation rate if we consider median temperature at 150 m in the Central Gulf to be the essential E variable capturing Greenland halibut production.

**Figure 8:**
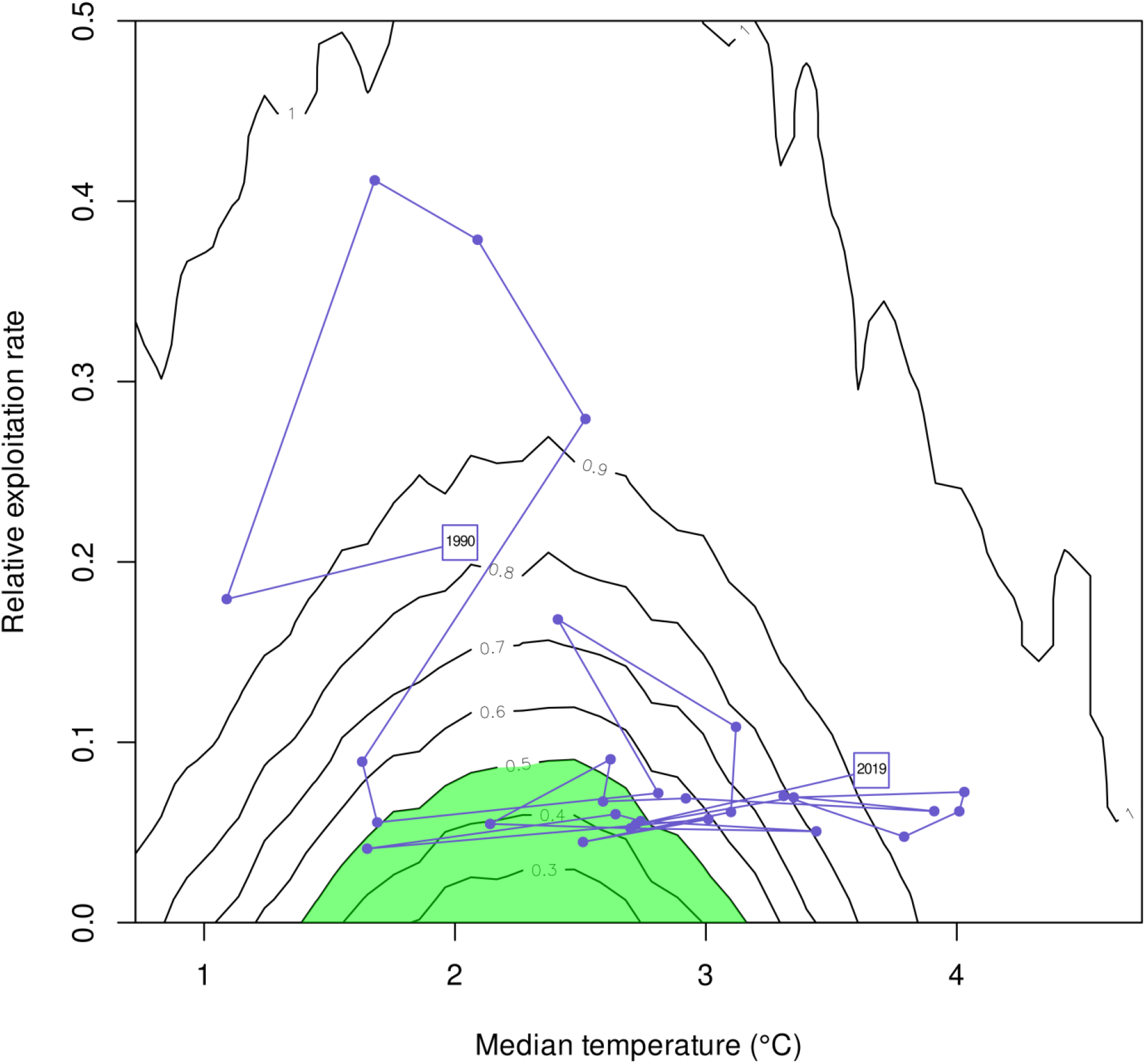
A contour plot showing how exploitation rate of the stock would need to be altered to condition advice for Gulf of St Lawrence turbot to changes in median bottom water temperature at 150 m depth. The climate impact is mediated through the relationship between the temperature and turbot production rate observed in the past. The contours represent the risk expressed as the probability of not achieving the target biomass objective (mean biomass from 1995-2000) after 10 years for different fishing and climate scenarios. For risk levels 50% and lower (shaded green), risk equivalent strategies are possible to compensate for CC by adjusting relative exploitation rate. The actual exploitation rate and temperature scenario is overlaid on the contours.

The track of the actual relative exploitation rate and the observed temperature (Fig 8, trace line) shows that even though the relative exploitation rates are not large for most demersal species (e.g. < 10%), the median temperature has already increased to well outside the area where Greenland halibut production is maximised. The observed 2019 temperature and the relative exploitation rate placed on the stock in 2019 places the risk to the stock of not achieving its objective in 10 years of fishing at this rate at >90%. The history of exploitation and temperature shown by the trace line suggests that the stock has often been outside the safe operating space but it has shown a systematic movement outside that space (in the temperature dimension rather than in exploitation) in the past several years. This suggests that the stock productivity state has shifted or continues to shift directionally to a poorer state. Thus, the contour lines can be used for showing how fishery removals have to be adjusted to maintain the desired probability of achieving a specified management objective and/or to provide risk equivalent advice on fishery exploitation conditioned on temperature (or “E” more generally), which is a surrogate variable for directional change (i.e. Climate Change conditioned advice).

## 5 Discussion

Decisions in the management of fisheries are always risk-based decisions, although both risk and fishery objectives may not always be explicit. A growing emphasis on transparency and rule-based management is increasing the use of both explicit objectives and explicit risk tolerances in fisheries management[32,33]. Managing fish stocks in a changing environment or ecosystem increases the value of risk analyses and a risk-based approach to management, if the risk quantification can capture added uncertainty due to the changing environment[34]. Because fisheries are already implicitly or explicitly managed in a risk-based framework, expressing the impacts of climate or ecosystem, change in terms of risk associated with fisheries decision making is fully consistent with the existing paradigm. The analytical approach proposed here focuses on the incremental uncertainty in stock production arising from a changing environment when quantifying the response of fish stocks to fishing pressure and the ability to achieve fishery management objectives, given a specific environment scenario. This uncertainty can be estimated regardless of the depth of detailed mechanistic understanding of stock productivity dependence on the environment even if the amount of that uncertainty might differ between approaches. Here we applied an EM method for developing climate-informed, risk-based science advice for a data-moderate fish stock in a rapidly changing environment. An EM approach allows us to take advantage of the available information and data to define a functional relationship between stock production and an environmental E variable, which can be modified and improved as more information becomes available in a manner consistent with a plausible hypothesis. Development of an EM model does not preclude the development of more detailed approaches later on that may capture uncertainty better, but they do allow us to proceed in the present rather than waiting for future developments before advice can be provided.

CC is accelerating[2] and there are indications of non-linearity and possibly tipping points in many features of the environment affected by CC[35,36], particularly in Canadian and other higher latitude waters[37]. The time scales of many of the projections in the IPCC reports are consistent with CC driven factors potentially affecting fish production within time frames where fisheries management advice on factors like target biomass and exploitation rates is required e.g. 10 years in the turbot case study explored here. Many jurisdictions have requirements for setting of fishery objectives (e.g. target and limit reference points for biomass and exploitation rate) or established recovery targets and plans for collapsed stocks embedded in legislation or binding policy. The ability to comply with such targets, limits and/or rebuilding plans can be affected by non-stationary environmental conditions that are in turn affected by CC. These types of requirements and management standards increase the need to provide fisheries advice that considers CC impacts as part of the body of advice for fishery managers. Developing this advice in a risk-based framework that informs decisions about risk equivalent exploitation is a particularly direct and transparent way for science to inform decision makers of the risk implications of their choices. Such advice has the strengths of being evidence-based, uses as little or as much information as is available about both changes in the environment of the stock and the impacts of environmental conditions on stock productivity, and clearly articulates the CC-related risks to the resource under various management options as risk increments (positive or negative) from status quo risk when CC is not considered (and the environment is assumed to be stationary).

Development of EMs is an effective way to condition fishery advice to CC in the short term to medium term. These models are relatively quick to implement and they tend to be robust because they are driven by broad phenomenological relationships and take advantage of emergent phenomena and constraints present in the climate forcing feedback system[38]. The EM approach should not be seen as contrary to detailed mechanistic approaches but complementary. EM methods can incorporate process-based understanding as knowledge increases, and when knowledge is sufficient, mechanistic approaches can take many of the elements of risk-based approaches from EM. Complementary approaches also provide consistency checks on one another in the same way that they have done for sea level rise predictions from global climate models[12]. The multiple model paradigm is a way to provide more robust risk analyses and overall science advice and quickly provide a means of conditioning advice to the CC reality.

When developing EMs for CC impacts on fish stock it is necessary to think clearly on what climate-driven environmental (E) variable to use and define a hypothesis for how fish stock production should change with E. Good choices will be case specific and require appropriate experts and/or resource users familiar with the stock and environment and the amount of information available, to make a wise choice of E. Some of the necessary characteristics of an appropriate E variable is that it should be hypothesised or demonstrated to have some impact on stock production and therefore could affect the achievability of stock objectives. The E variable should also have a hypothesized or demonstrated link to how climate change is affecting the general area, or to some other environmental forcing factor considered by experts to be potentially important to changes in the marine ecosystem. For the Greenland halibut in the Gulf of St. Lawrence, we selected water temperature in the central Gulf at 150 m depth as the E variable. This area is not the main region where Greenland halibut is concentrated and fished in the Gulf (Fig. 1); however, northward flowing water in the Gulf branches at the central Gulf and feeds both the main eastern and western turbot fishing areas[39]. Therefore, the central Gulf temperature does have bearing on all the main turbot fishing areas and therefore it is a measure that is likely to have a bearing on what the fish experience. This choice of E variable was also based on considerations of explanatory power (AIC value) and the expectation of a mono-modal-type relationship between turbot production and temperature since the species is adapted to a relatively narrow range of temperature conditions[40] (S2 Appendix). E variable selection may take into account consequences of a variety of processes including physiological and ecological adaptation, it may consider time lags, and it may relate to other driving forces on stock production such as recruitment or mortality. Therefore, it is important to consider that E may not be a direct driver but could potentially behave as a meta-variable pertinent to the productivity of a particular stock. The shape of the production vs E relationship should also be considered. The E variable here showed a dome shaped relationship with E (Fig. 3) which is the logical shape one might expect for a population adapted to a relatively narrow range of conditions where its productivity decreases outside that range. Nonparametric approaches model fitting approaches like GAMs can be used to try to quantify empirical relationships that may be present. GAMs are particularlyHowever, there must be an ability to infer plausible interpretations of relationships that are found, and if possible, suggest more targeted analyses or experiments to improve understanding of the bases for the relationships useful in these tasks because they develop a central tendency for the relationship within interpolated ranges while still allowing considerable freedom in the form of the P/B vs E relationship. When information is adequate, one can take into account more than one E variable and their combined impact on the P/B ratio of the stock. Multivariate GAMs are also easily fitted if there is a sufficient amount of data. However, as the number of E variables increases, plausible future scenarios are needed for each one and the potential interactions and non-linearities among scenarios for different E variables must be considered. Thus, we recommend the KISS (keep it simple and straightforward) principle in choosing E variables for this empirical approach, and interpreting E variables as capturing an ensemble of environmental forcing processes rather than very specific mechanisms.

An alternative approach to projecting the stock into the future is to sample the actual surplus production values from certain time periods where E conditions had different means, i.e. block sampling or by sampling from particular quantiles of the E distribution. This does free one from forming a hypothesis on the mechanism, which though elegant, can only produce values that have been observed previously and is constraining especially for shorter time series. This block resampling approach limits the range of values of future E in scenarios and the precedents being set in many regions of the world with environmental variables such as temperature with every new year of data suggests that this is likely to be a conservative approach that may well underestimate the impact of climate change on stock production. And although that approach may appear hypothesis-free, it is still making a strong assumption about future production being the same as some part of the past. We also advocate the approach that forces an explicit hypothesis on P/B vs E because it is more clearly falsifiable.It is constraining because it

Density dependence is a key consideration in fisheries models so that stocks do not grow exponentially beyond natural limits. The EM approach used here includes a simple form of density dependence by introducing a carrying capacity parameter (eq. 8) in the same manner as the Schaefer production model[13], i.e. it acts as a reducing factor in growth rate to the point where growth is zero when biomass is at the carrying capacity. The carrying capacity parameter was set at three times the maximum observed biomass since the survey started in 1990, which is arbitrary but does take into consideration that the stock has been fished commercially for >50 years[41] and may once have been much larger than biomasses estimated from the survey series. Density dependence did not have an appreciable impact on model outputs, nor on estimates of the incremental risk associated with the inclusion of CC in stock projections. This lack of influence of density dependence on projections from current stock sizes is a useful result because it means for stocks assessed as well below their carrying capacities, CC conditioning with EMs can be done without including density dependence parameters in the EM projections. For stocks that are in very depressed states where depensation is considered a plausible hypothesis[42], it may be necessary to include a depensatory process in the model, and for projections that suggest stocks may return to much larger sizes in just one or two decades, monitoring for evidence of density dependence in stock dynamics can be implemented as the stock grows.

## 6 Summary

We developed an EM approach that describes how fish production changes as a function of a CC variable and carries that relationship through to a risk assessment and estimation of the incremental risk for alternative fishing strategies in a context of CC. Using basic time series data that are typically available for exploited demersal fish stocks, the model was parameterised and projected forward, given future climate scenarios inferred from a single environmental variable. The risk assessment required an explicit management objective, which in this case corresponded to a target objective defined by setting a baseline period over which to average stock status. Model projections under different climate scenarios could then be compared to a baseline or ‘average conditions’ climate scenario. The comparisons formed the basis for determining the incremental risk associated with each scenario under a status quo fishing strategy (assuming random or constant environment with no productivity dependence on the environmental variable), and the change in fishery exploitation rates required to maintain the risk at a specified tolerance - in this case 50% likelihood of achieving the objective (i.e., risk equivalent fishery exploitation advice). This approach shows how fishing pressure should change in relation to CC (expressed as E) if the management goal is to achieve the stock objective with the same level of risk. It can also inform managers on the relative change in risk associated with alternative management options under different climate scenarios, fostering transparency into decisions made considering CC-contributed uncertainty. This approach can therefore use previously observed stock productivity dynamics to define a safe operating space for fisheries given the projected future climate, the management objective and timeframe and the risk level considered acceptable. The approach can be used to expand stock exploitation advice to take CC into account, inform managers of the change in risk that can be anticipated under different climate scenarios, and support a transparent dialogue among scientists, managers, and stakeholders on how risk decisions can be altered by CC, by giving a rough magnitude of the changes potentially required in stock exploitation advice if CC is to be considered.

Although this risk-based approach was designed to develop advice that considers CC, it can be applied to any ecosystem or environmental driver affecting stock production, for example in cases of habitat loss not caused by CC, or where there is potential for a degree of interdependence between sustainable exploitation rates for a predator and a prey species. The main condition to applying this risk-based approach to other potential drivers is the need for a plausible hypothesis on how the external variable affects a stock’s production. At an even higher level than fish stock production, the risk equivalent principles for altering advice are more universal and can work in any area of natural resource management where status quo advice is considered possibly suspect because of a change in an external factor

## 7 Acknowledgements

Johanne Gauthier, Peter Galbraith and Denis Gilbert for providing the stock, temperature and oxygen data, respectively. Important feedback on many of the ideas presented here was provided by participants of the DFO peer review meeting in incorporating CC into stock assessment advice in May 2018. Mariano Koen-Alonso suggested that density dependent processes should be explored.

